# Glutamine is required for M1-like polarization in response to *Mycobacterium tuberculosis* infection

**DOI:** 10.1101/2022.01.11.475775

**Authors:** Qingkui Jiang, Yunping Qiu, Irwin J. Kurland, Karl Drlica, Selvakumar Subbian, Sanjay Tyagi, Lanbo Shi

## Abstract

In response to *Mycobacterium tuberculosis* infection, macrophages mount early proinflammatory and antimicrobial responses similar to those observed in M1 macrophages classically activated by LPS and IFN-γ. A metabolic reprogramming to HIF-1-mediated uptake of glucose and its metabolism by glycolysis is required for M1-like polarization, but little is known about other metabolic programs driving M1-like polarization during *M. tuberculosis* infection. Identification and quantification of labeling patterns of U^13^C glutamine and U^13^C glucose-derived metabolites demonstrated that glutamine, rather than glucose, is catabolized in both the oxidative and reductive TCA cycle of M1-like macrophages, thereby generating signaling molecules that include succinate, biosynthetic precursors such as aspartate, and the antimicrobial metabolite itaconate. This conclusion is corroborated by diminished M1 polarization via chemical inhibition of glutaminase (GLS), the key enzyme of the glutaminolysis pathway, and by genetic deletion of *GLS* in infected macrophages. Furthermore, characterization of the labeling distribution pattern of U^15^N glutamine in M1-like macrophages indicates that glutamine serves as a nitrogen source for the synthesis of intermediates of purine and pyrimidine metabolism plus amino acids including aspartate. Thus, the catabolism of glutamine, as an integral component of metabolic reprogramming in activating macrophages, fulfills the cellular demand for bioenergetic and biosynthetic precursors of M1-like macrophages. Knowledge of these new immunometabolic features of M1-like macrophages is expected to advance the development of host-directed therapies that will enhance bacterial clearance and prevent immunopathology during tuberculosis.

**Summary:** Recent advances in immunometabolism have stimulated increasing interest in understanding the specific cellular metabolic states of immune cells associated with the various disease states of tuberculosis. As the primary target of *Mycobacterium tuberculosis* (*Mtb*) infection, macrophages play essential roles in dictating the progression and final outcome of infection. Previous studies, including our own, show that the upregulation of hypoxia-inducible-factor 1 alpha (HIF-1α) and a metabolic reprogramming to the Warburg effect-like state are general features of the host immune cell response to *Mtb* infection. They are also critical for macrophage polarization to the M1-like phenotype characterized by high-level expression of proinflammatory and antimicrobial molecules against *Mtb* infection. However, our knowledge about the immunometabolic features of M1-like macrophages is poor. Using widely targeted small metabolite (WTSM) screening (600+ small polar metabolites) and stable isotope tracing of U^13^ glutamine, U^13^ glucose, and N^15^ glutamine, as well as therapeutic and genetic approaches, we report that, in addition to elevated glucose catabolism by glycolysis, glutamine serves as an important carbon and nitrogen source for the generation of building blocks, signaling molecules, and antimicrobial metabolite during macrophage polarization to the M1-like phenotype. The study adds novel insights into the immunometabolic properties of *Mtb*-infected macrophages.

## Introduction

As professional phagocytes, macrophages play essential roles in regulating tissue homeostasis and immune response to pathogens. Although traditionally classified into proinflammatory M1 and anti-inflammatory M2 states by their response to treatment with IFN-γ and LPS, as well as with IL-4 and IL-13, respectively (1), the states of macrophage activation are often within the spectrum of M1 to M2, depending on microenvironmental factors and signaling molecules (2, 3). Nevertheless, recent advances in immunometabolism reveal that the polarization states of macrophages are closely associated with distinctive metabolic states. M1 polarization is marked by increased expression of glucose-uptake transporters and isoenzymes of the glycolysis pathway, which leads to elevated glycolytic flux with increased lactate formation and secretion (4), similar to the Warburg Effect (aerobic glycolysis) seen in cancer cells. In contrast, M2 polarization depends predominantly on mitochondrial oxidative metabolism (4, 5).

Amino acids also play important roles in macrophage activation (6). For example, it is well known that increased uptake of arginine and its catabolism by inducible nitric oxide synthase 2 (NOS2) produces NO, which is indispensable for the defense of M1 macrophages from invading pathogens. In contrast, arginase 1-mediated metabolism supports M2 polarization, tissue homeostasis, and repair (1). Additionally, catabolism of tryptophan by amino acid oxidases, including IL4I1 and/or indoleamine 2,3-dioxygenase 1 with the formation of bioactive metabolites, such as kynurenine and kynurenic acid, promotes the generation of repressor macrophages and inhibition of Th1 immunity (7–10). Glutamine, which is consumed by immune cells at a similar or higher rate than glucose, is an essential nutrient for effector functions of activated immune cells (11). However, a clear role for glutamine in macrophage polarization has not been established due to conflicting reports. For example, glutamine is proposed to be important for signaling by hypoxia-inducible factor 1 (HIF-1) and for mTOR1C activation in LPS-induced M1 macrophages (12, 13). In contrast, several other studies argue that glutamine metabolism restricts the proinflammatory M1 state and favors M2 polarization (14–16). The latter assertion derives from, at least in part, the glutamine-derived alpha ketoglutarate (α-KG) that enhances mitochondrial oxidative metabolism and inhibits HIF-1α by promoting the activity of HIF prolyl hydroxylases (14). Thus, the role of glutamine during infection with *Mycobacterium tuberculosis* (*Mtb*) needs to be established.

Macrophage infection by various bacteria, including *Mtb*, the etiological agent of tuberculosis (TB), leads to an initial robust proinflammatory response that resembles the classically activated M1 macrophages (17, 18). Previous studies, including our own, reveal a metabolic reprogramming involving HIF-1 induction and increased glycolytic flux, indicating a role of glycolysis pathway in the proinflammatory response of *Mtb*-infected murine bone marrow-derived macrophages (BMDMs) (19–22). A similar metabolic reprogramming is also required to activate human macrophages in response to *Mtb* infection, as perturbation of glycolysis by 2-deoxyglucose dampens the M1-like polarization and promotes survival of *Mtb* (23). To date, the other metabolic programs propelling the M1-like polarization during *Mtb* infection have not been clearly defined. Given that *Mtb* can modulate the immunometabolic response of infected macrophages to survive and persist (24), a better understanding of metabolic programs of host cells will help develop host-directed therapies (HDTs) to promote bacterial clearance.

In the present work we dissected the immunometabolic features of *Mtb* infection-induced M1-like macrophages using 1) metabolic profiling and stable isotope-assisted metabolomics, 2) single-molecule RNA fluorescence *in situ* hybridization (sm-RNA-FISH), and 3) therapeutic or genetic manipulations targeting glutaminolysis. We report that, apart from increased glucose catabolism by glycolysis, glutamine and its direct metabolite, glutamate, serve as carbon and nitrogen source for the synthesis of biosynthetic precursors involved in multiple metabolic pathways of activating macrophages. Thus, glutamine is central to the proinflammatory and antimicrobial responses of M1-like polarization against *Mtb*.

## Results and Discussion

### *Characterization of infected bone marrow-derived macrophages* by sm-RNA-FISH

Macrophage activation during *Mtb* infection shows an early M1-like phenotype accompanied by a metabolic shift to glycolysis for at least 8 to 12 hrs post infection (p.i) (22, 25). That is followed by a late resolution/adaptation phase associated with dampening of M1 polarization, and a recovery of mitochondrial oxidative metabolism at 24 hrs p.i and beyond (21, 25). Such a shift in the immunometabolic states of infected macrophages corresponds to changes in growth dynamics and physiology of *Mtb* inside host cells (26). We used sm-RNA-FISH, which allows specific detection and quantification of single molecules of target mRNAs *in situ* at a single-cell level (27), to characterize the expression of several immunometabolic markers of infected murine BMDMs. Consistent with the transcriptomic dynamics of *Mtb*-infected BMDMs (28, 29), infected BMDMs at 4 to 8 hrs p.i showed increased mRNA molecules of the M1 markers IL-1β and NOS2, as well as the Warburg Effect enzymes that include GLUT1 (glucose transporter 1), PFKFB3 (6-phosphofructo-2-kinase/fructose-2,6-bisphosphatase 3, a key regulatory enzyme for glycolysis), and MCT4 (the major lactate efflux transporter) (**Fig. 1A-F**). These increases were followed by a decrease at 24 hrs p.i. In contrast, mRNA molecules for ARG1, an M2 marker, showed only a clear induction at 24 hrs p.i (**Fig. 1G**). High-level heterogeneity was observed among the cells, with only some cells expressing a significant number of mRNA molecules for a given marker (**Fig. 1A**). This observation is consistent with the stochastic nature of mRNA synthesis (30, 31). The mRNA expression dynamics of the immunometabolic markers correlated with the production of IL-1β by infected BMDMs, as measured in the culture supernatant (**S. Fig. 1**). Thus, murine BMDM infection was a suitable model system for studying the metabolic programs of M1-like macrophages.

**Fig. 1.**
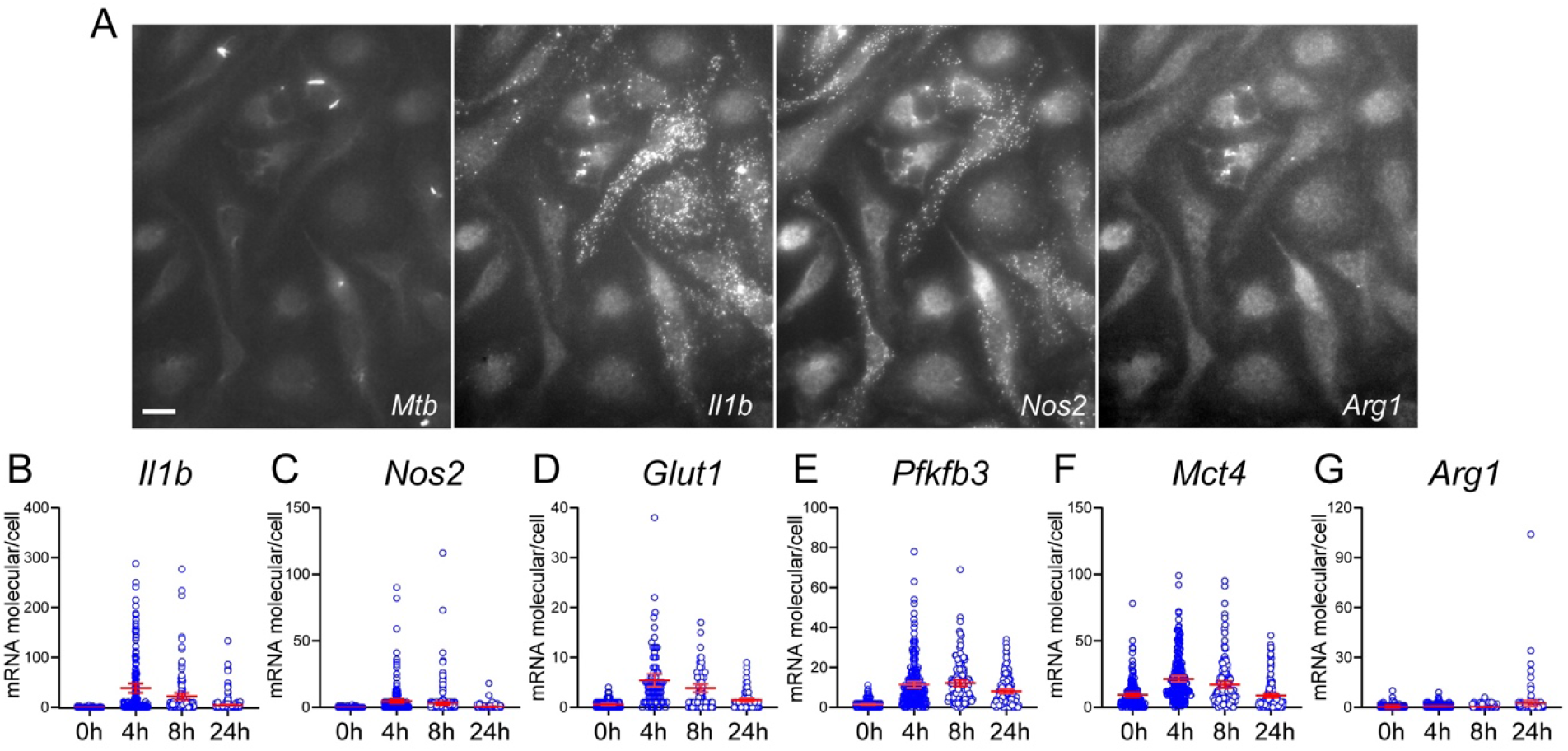
Single-cell mRNA analysis of immunometabolic markers in *Mtb*-infected BMDMs. BMDMs seeded on coverslips were probed in 3-plex hybridization reactions using sm-RNA-FISH probes labeled with transcript-specific probe sets (about 50 oligonucleotides for each mRNA) that were coupled to tetramethylrhodamine (TMR), Texas Red, or Cy5 fluorophores. Images were acquired in Z-stacks of different fields of cells in different channels. Fluorescence spots corresponding to single mRNA molecules in individual cells were counted in the merged Z-stacks using a custom image-processing algorithm implemented in MATLAB. Representative images of GFP-labeled *Mtb* and mRNA molecule spots for *Il1b* (TMR)*, Nos2* (Texas Red), and *Arg1* (Cy5) at 8 hrs p.i in Z-stacks of the same field of cells in different channels (**A**). Changes of respective mRNA molecules of immunometabolic markers in individual cells at hr 0, 4, 8, and 24 p.i. (**B - G**). A total of ~ 100 - 150 cells were analyzed by sm-RNA-FISH at indicated times p.i. Each circle represents one cell. Representative data are shown as means ± 95% CI (Confidence Interval) from at least three independent experiments. The scale bar in A is 10 μm.

### Metabolite screening identifies highly impacted metabolic pathways associated with glutamine

To better understand the metabolic programs driving the M1-like polarization, we carried out a widely targeted screen of small metabolites by the QTRAP 6500+ LC-MS/MS Systems (ABSciex) in *Mtb*-infected BMDMs at 8 hrs p.i. A total of 169 metabolites were detected having a coefficient of variation (CV) below 30% in the multi-injection of quality-control samples. Multivariate analysis of the data sets was performed using SIMCA-p software; the partial least squares-discriminant analysis (PLS-DA) model revealed distinct differences in the metabolic profiles between infected macrophages and uninfected controls (**Fig. 2A**). Key metabolites that contributed to this distinction were selected by the variable importance in the projection (VIP > 1).

**Fig. 2.**
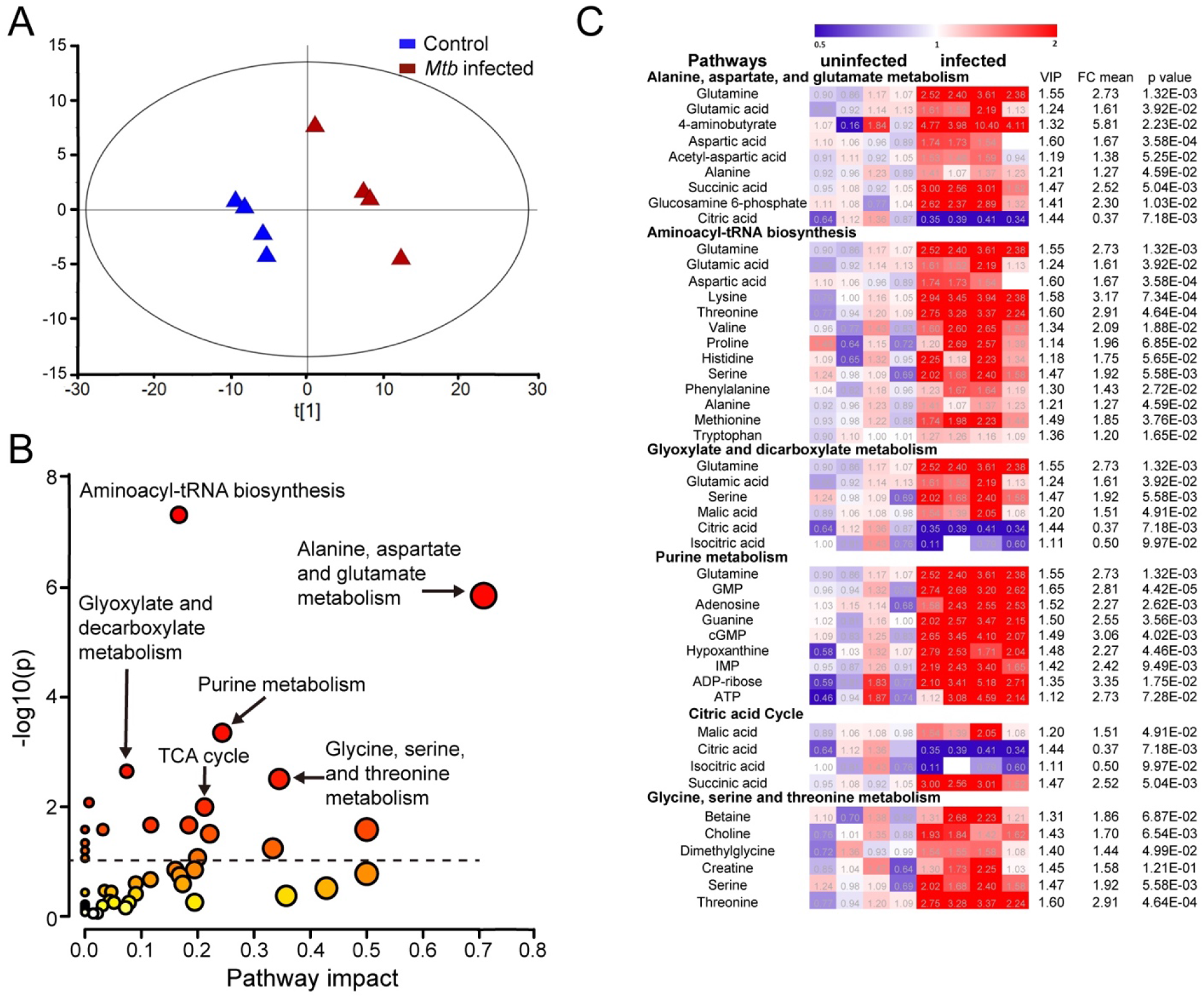
Identification of highly impacted metabolic pathways during *Mtb*-induced M1-like polarization by widely targeted small-metabolite screening. Metabolites extracted from *Mtb*-infected BMDMs at 8 hrs p.i and uninfected controls were analyzed by the QTRAP 6500+ LC-MS/MS systems. Separation of M1-like macrophages from uninfected controls as scores plot from PLS-DA modeling (**A**). Highly impacted metabolic pathways in M1-like macrophages (**B**). The differential metabolites with variable importance in the projection (VIP) > 1 from the PLS-DA modeling of the two groups were subjected to pathway enrichment analysis by the Metaboanalyst (V5.0). The circle size denotes the altitude of pathway impact, and color darkness represents the extent of significance. Heatmaps of differential metabolites with VIP > 1 in highly impacted metabolic pathways (**C**). Data are shown as normalized values to the corresponding mean value of the uninfected group.

As expected, increased levels of glycolytic intermediates, such as fructose 1,6-bisphosphate and 3-phosphoglyceric acid, in infected BMDMs (**S. Fig. 2**) were consistent with increased glycolysis during the macrophage response to *Mtb* infection (19, 22, 23). Using pathway enrichment analysis (Metaboanalyst (V5.0)) with the differential metabolites that were selected based on PLS-DA analysis, we identified highly impacted metabolic pathways associated with macrophage activation to the M1-like phenotype (**Fig. 2B**). A detailed examination of metabolites among the impacted pathways revealed that glutamine and its direct metabolite, glutamate, had positive associations with multiple pathways of macrophage activation, including purine metabolism, glyoxylate dicarboxylate metabolism, the TCA cycle, as well as alanine, aspartate, and glutamate metabolism. The latter was the most affected metabolic pathway, having a low - log10(p) of 6 and high pathway impact factor of ~ 0.7 (**Fig. 2B-C**).

Strong evidence for glutamine involvement in M1-like polarization was the identification of increased accumulation in infected BMDMs of 4-aminobutyrate (GABA), which could be derived from the GABA shunt that supplies glutamate-derived succinate to the TCA cycle. The GABA shunt is associated with M1 polarization, as inhibition of the shunt by vigabatrin (an irreversible inhibitor of GABA transaminase) decreases succinate concentration, leading to reduction of HIF-1 and IL-1β in LPS-activated M1 macrophages (13). The identification of increased 2-hydroxyglutaric acid (2HG) (**S. Fig. 2**), which is converted from glutamine-derived α-KG in cancer cells having deficient activity of isocitrate dehydrogenase 1 or 2 due to mutation, and in mammalian cells under hypoxia (32–36), also suggests the connection of glutamine catabolism with the TCA cycle during the M1-like polarization. Given that glutamate participates in the synthesis of glutathione (GSH), a major intracellular, small-molecule antioxidant in proinflammatory immune cells (37, 38), increased levels of GSH in both the reduced and oxidized forms (**S. Fig. 2**) supports a role for glutamine metabolism in maintaining redox homeostasis of M1-like macrophages. Collectively, the findings from metabolite screening of infected macrophages suggest an important role for glutamine catabolism in M-1 like polarization.

### Tracing metabolomics of U^13^C glutamine and glucose identifies anaplerosis of glutamine carbon through both the oxidative and reductive TCA cycle

To determine the contribution of glutamine as a carbon source for the metabolic program of macrophage activation, we used U^13^C glutamine to track its metabolic signature in *Mtb*-infected M1-like macrophages by Gas Chromatography/Time-of-Flight Mass Spectrometry (GS-TOF/MS) (39). Infected BMDMs were cultured for 8 hrs in DMEM medium supplemented with 4 mM 50% U^13^C glutamine; then cellular metabolites were extracted, derivatized with silylation reagents, and analyzed for isotope enrichment (39). Carbons of glutamine can enter the TCA cycle in the form of glutamate-derived α-KG, and/or as succinate from the GABA shunt (**Fig. 3A**). The labeling pattern of U^13^C glutamine carbons from the first turn of the oxidative TCA cycle would generate four ^13^C labeled intermediates (detected as M+4 by mass spectroscopy), such as succinate, fumarate, malate, and citrate. The labeling pattern of U^13^C glutamine from the first turn of the reductive TCA cycle would produce M+5 intermediates citrate and isocitrate (**Fig. 3A**). As shown in **Fig. 3C - I & S. Table 1**, increased enrichment of ^13^C isotopes was predominantly for M+4 intermediates (~ 8 – 12%), including succinate, fumarate, malate, itaconate, and citrate, as well as for M+5 citrate (~ 13%) in infected BMDMs relative to uninfected controls. These data indicate that increased glutamine carbon flux to the TCA cycle intermediates arises from both the oxidative and reductive directions. This conclusion is supported by the simultaneous increase in enrichment for both M+4 (~ 1.6 fold) and M+5 citrate (~ 3.3 fold) in infected BMDMs (**S. Table 1**), with the latter deriving from the reductive carboxylation of glutamine-derived α-KG being more pronounced (**Fig. 3G**).

**Fig. 3.**
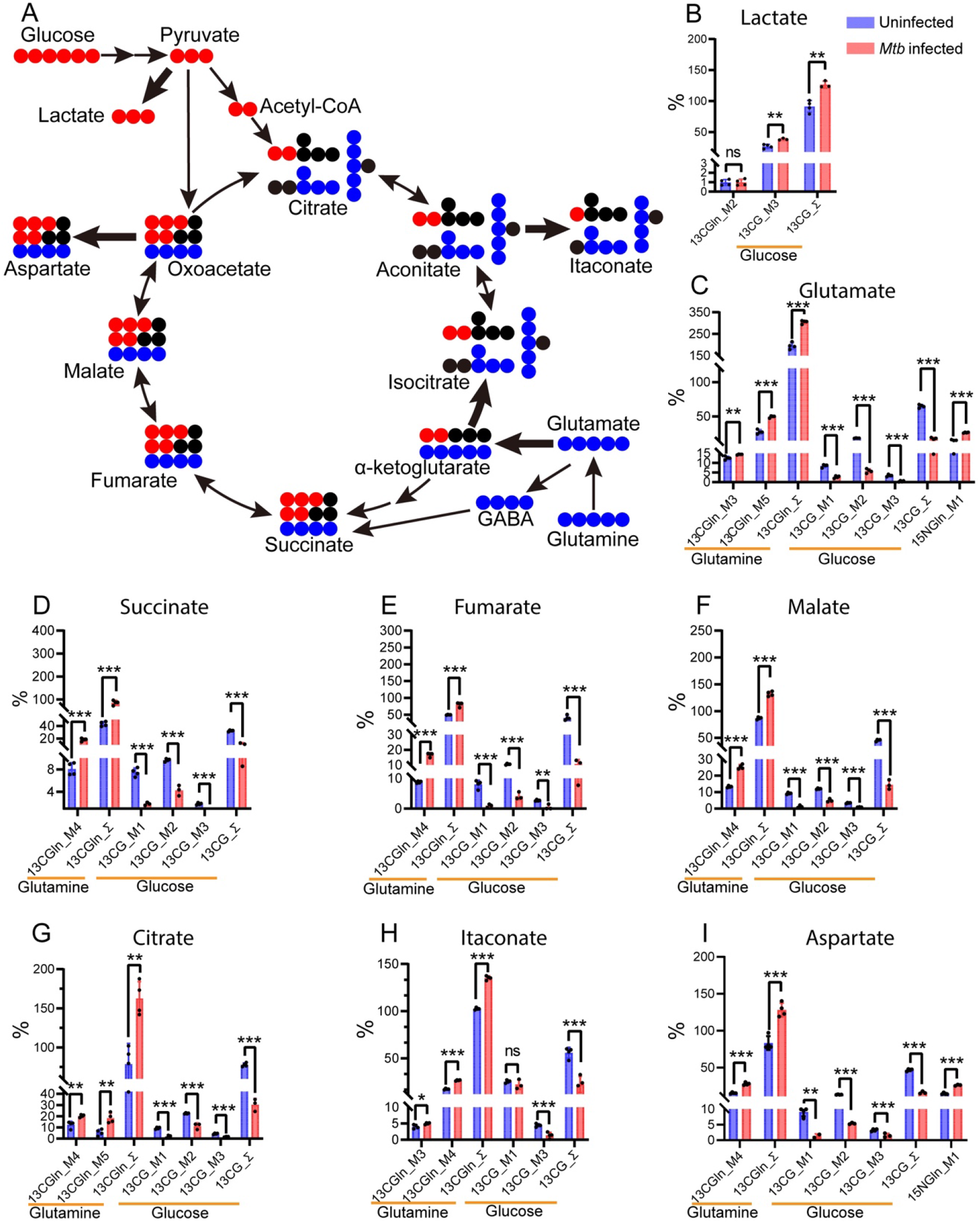
Isotope labeling distribution pattern of U^13^C glucose and glutamine, and U^15^N glutamine during the M1-like polarization. Diagram of ^13^C distribution from the catabolism of U^13^C glucose and U^13^C glutamine in glycolysis and/or the TCA cycle (**A**). Catabolism of U^13^C glucose generates M+3 glycolytic intermediates, and M+2 (via pyruvate oxidation) and M+3 (via pyruvate carboxylation) TCA cycle intermediates/derivatives. M+5 glutamate, the direct metabolite of U^13^C glutamine, can enter the TCA cycle in the form of alpha ketoglutarate (αKG) and/or via the GABA shunt. Increased ^13^C distribution from U^13^C glucose catabolism in glycolysis with the generation of M+3 lactate (**B**), and from the catabolism of U^13^C glutamine in the oxidative and reductive TCA cycle resulting in the generation of TCA cycle intermediates/derivatives (**C - I**). M1-like polarization was marked by diverting of the ^13^C glucose carbon distribution from the TCA cycle M+2, M+1, and M+3 intermediates to the formation of M+3 lactate (B). In contrast, catabolism of U^13^C glutamine led to increased ^13^C distribution in the form of M+4 TCA cycle intermediates/derivatives, including succinate (D), fumarate (E), malate (F), itaconate (H) and aspartate (I), as well as M+4 and M+5 citrate (G), indicating the simultaneous operation of both the oxidative and reductive TCA cycle. Increased ^15^N distribution from U^15^N glutamine to M+1 aspartate (I) also indicates glutamine being a nitrogen source for the formation of nonessential amino acid aspartate. Solid red and blue circles represent ^13^C from U^13^C glucose and U^13^C glutamine, respectively. Thick arrows indicate increased ^13^C carbon flux into the formation of lactate and the TCA cycle intermediates or derivatives, respectively. Enrichment calculated from U^13^C glucose was marked as 13CG_M1, 13CG_M2, 13CG_M3, and 13CG_∑. ∑ was calculated with M1*1+M2*2…+Mn*n. Enrichment calculated from U^13^C glutamine was marked as 13CGln_M3, 13CGln_M4, 13CGln_M5, and 13CGln_∑. Enrichment calculated from ^15^N glutamine was marked as 15NGln_M1. Data are shown as means ± SD from 3 - 4 biological replicates. Statistical significance at \**P* < 0.05, \*\**P* < 0.01, \*\*\**P* < 0.001 and \*\*\*\**P* < 0.0001 was based on two-tailed Student’s T-test.

The decrease of citrate and isocitrate in infected BMDMs (Fig. 2C) and increased enrichment of M+4 and M+5 citrate from U^13^C glutamine in both the oxidative and the reductive directions of the TCA cycle suggest an outflow of mitochondrial citrate. Indeed, the ~10-fold increase in the ratios between total itaconate and total citrate in infected BMDMs (**S. Fig. 3**) is consistent with the high-level induction of the aconitate carboxylase-1 gene (*Acod1* or *Irg1*), whose product catalyzes the formation of itaconate in M1 macrophages (22, 28, 40, 41). This conclusion is supported by the increased enrichment for M+4 itaconate (**Fig. 3H**), which could be derived from the oxidative TCA cycle-generated M+4 and/or the reductive TCA cycle-generated M+5 citrate. The enhanced formation of itaconate, and its inhibition of succinate dehydrogenase activity (42), may be a driving force in directing the flux of glutamine toward succinate, as recently reported (43).

Interestingly, we also observed very low enrichment of M+3 TCA cycle intermediates/derivatives in M-1 like macrophages (**S. Table 1**). These data indicate that the reductive TCA cycle-derived M+5 citrate is not utilized for *de novo* fatty acid synthesis through the cytosolic citrate lyase-mediated pathway, but rather it is redirected to the formation of itaconate in M1-like macrophages. This finding contrasts with the utilization of M+5 citrate for fatty acid synthesis, which consequently leads to the generation of M+3 TCA cycle intermediates, such as malate, fumarate, and succinate, in cancer cells under hypoxia and impaired respiration (44, 45). Additionally, the absence of M+3 lactate in M1-like macrophages (**S. Table 1**) indicates glutamine is not a significant carbon contributor for lactate formation. Importantly, the increased enrichment of ^13^C for M+4 aspartate (~ 1.9 fold) in infected BMDMs (**Fig. 3I & S. Table 1**), which probably derives from transamination reactions of the TCA cycle intermediate oxaloacetate (OAA) (46), also shows glutamine to be the major carbon donor for *de novo* synthesis of the nonessential amino acid aspartate during M1-like polarization. Taken together, the U^13^C glutamine tracing metabolomics data indicate that during M-1-like polarization, glutamine replenishes TCA cycle carbon flux for the synthesis of signaling molecules, such as succinate, antimicrobial metabolite itaconate, and nonessential amino acids such as aspartate.

To confirm that glutamine rather than glucose serves as a major carbon source for the TCA cycle during M1-like polarization, we analyzed the metabolic signature of U^13^C glucose by GS-TOF/MS(39). Infected BMDMs were cultured for 8 hrs in DMEM medium supplemented with 25 mM of 50% U^13^C glucose, and cellular metabolites were extracted for isotope enrichment analysis. The labeling pattern of U^13^C glucose from glycolysis and the first turn of the TCA cycle would result in the generation of M+3 glycolytic intermediates plus M+2 and M+3 TCA cycle intermediates from pyruvate dehydrogenase (PDH)- and pyruvate carboxylase (PC)-mediated pathways, respectively (**Fig. 3A**). As expected, the increased enrichment of ^13^C M+3 lactate (~ 12%) in *Mtb*-infected BMDMs, relative to uninfected controls (**Fig. 3B**), agreed with increased glycolytic flux from glucose to lactate during the M1-like polarization (19, 23). Importantly, in contrast to the labeling distribution of U^13^C glucose in uninfected controls, which is marked with ^13^C isotope distribution such as M+2, M+3, and M+1 (deriving from the second turn of the PDH pathway) into TCA intermediates/derivatives that include glutamate (**Fig. 3C - I**), M1-like polarization during *Mtb* infection was marked by decreased enrichment of ^13^C isotopes for the TCA-cycle intermediates, as well as their sum (∑_m_^n^) (**Fig. 3D - I & S. Table 1**), including succinate (~ 2.9 fold), fumarate (~ 3.5 fold), malate (~ 3.1 fold), and citrate (~ 2.6 fold) plus their derivatives itaconate (~ 2.1 fold) and aspartate (~ 2.7 fold).

The results described above support our previous work that shows diminished PDH function in *Mtb*-infected macrophages (22). In addition, given that itaconate is derived from M+2 citrate precursor, our observation of little change in M+1 itaconate (**Fig. 3H**), compared with a 2-fold decrease in M+2 citrate (**Fig. 3G**) in infected M1-like macrophages, indicates a high rate of conversion of newly synthesized citrate toward the formation of itaconate in infected macrophages. This result is consistent with high-level induction of *Irg1* in infected macrophages (25, 41), despite the decrease in overall flux to itaconate from glucose, as discussed above. Taken together, these data demonstrate the rerouting of ^13^C glucose carbon flux from the TCA cycle to glycolysis during M1-like polarization. This rerouting is consistent with the presence of the Warburg Effect in *Mtb*-infected macrophages (19, 22, 23) and with the observation that glucose is not a major carbon donor for the TCA cycle and aspartate synthesis in M1 macrophages (47).

### Tracing metabolomics of U^15^N glutamine identifies glutamine as nitrogen source for the synthesis of nucleotides and amino acids

By using U^15^N glutamine as a possible nitrogen source, we tracked the incorporation of glutamine nitrogen in the synthesis of nitrogen-containing compounds in M1-like macrophages by GC-TOF/MS (39). Infected BMDMs were cultured for 8 hrs in DMEM medium supplemented with 4 mM of 50% U^15^N glutamine, and cellular metabolites were extracted for isotope enrichment analysis. Consistent with increased purine metabolism during M1-like polarization, as seen in the unlabeled metabolomics studies discussed above, we found increased enrichment of ^15^N glutamine in key metabolites of purine metabolism, including hypoxanthine (M+1: ~ 4.7 fold; M+2: ~ 2.5 fold), adenine (~ 1.5 fold for both M+1 and M+2) and uric acids (~ 2.5 fold for both M+1 and M+2) (**S**. **Table 1**).

Since the generation of hypoxanthine and adenine is associated with nucleotide salvage pathways (47), the enrichment of glutamine nitrogen in their formation suggests a role for glutamine as a nitrogen source in the synthesis of nucleotides during M1-like polarization. In addition, enrichment of M+1 and M+2 uric acid is consistent with the finding that its formation via xanthine oxidase arising from the catabolism of hypoxanthine and/or xanthine is required for M1 polarization through xanthine oxidase-mediated ROS production and signaling (48). Increased enrichment for M+1 (~ 1.8 fold) and M+2 (~ 2.7 fold) uracil, which can be recycled by uridine phosphorylase or uracil phosphoribosyltransferase, also indicates an involvement of glutamine in the pyrimidine metabolism of M1-like macrophages.

Glutamine nitrogen was also participated in the synthesis of nonessential amino acids. M+1 aspartate and M+1 alanine were ~ 1.7 and 2.0-fold higher in M1-like macrophages, respectively, compared with uninfected controls (**Fig. 3I** & **S. Table 1**). It is worth noting that besides being a major carbon donor for the synthesis of aspartate, glutamine supplies reduced nitrogen to aspartate, probably as a result of mitochondrial transamination of glutamine-derived oxaloacetate in the TCA cycle (46). Glutamine-derived aspartate could contribute to the purine nucleotide cycle, which helps maintain the balance of the glycolysis-mitochondrial redox interface by preventing cytoplasmic acidification of M1 macrophages (47). This notion is supported by the recently identified role of FAMIN (Fatty Acid Metabolism-Immunity Nexus) as a multifunctional purine nucleoside enzyme activity that enables the purine nucleotide cycle (47). Indeed, expression of genes encoding purine-nucleotide cycle enzymes, including FAMIN, adenylosuccinate lyase, and AMP deaminase 3, are increased in *Mtb*-infected M1-like macrophages (28, 49). These data clearly demonstrate that glutamine serves as a nitrogen source for the metabolic reprogramming of M1-like macrophages.

### Chemical inhibition and genetic manipulation of glutaminolysis pathway diminishes M1-like polarization

To corroborate findings from metabolomics studies, we investigated the kinetics of glutamine uptake in infected BMDMs by monitoring changes of glutamine levels in the culture supernatant. We found that a high rate of glutamine uptake at early phases of macrophage infection (up to 8 hrs p.i) (**Fig. 4A**) coincided with M1-like polarization (refer to Fig. 1) and with increased mRNA molecules of *Gls* gene (**Fig. 4B**), which encodes the mitochondrial glutaminase (GLS), a key enzyme of the glutaminolysis pathway. An increased dependence on glutamine in *Mtb*-infected human monocyte-derived macrophages and THP1 cells at 18 hrs p.i is reported, although it is not clear about the contribution of glutamine to M1 and/or M2 polarization (50).

**Fig. 4.**
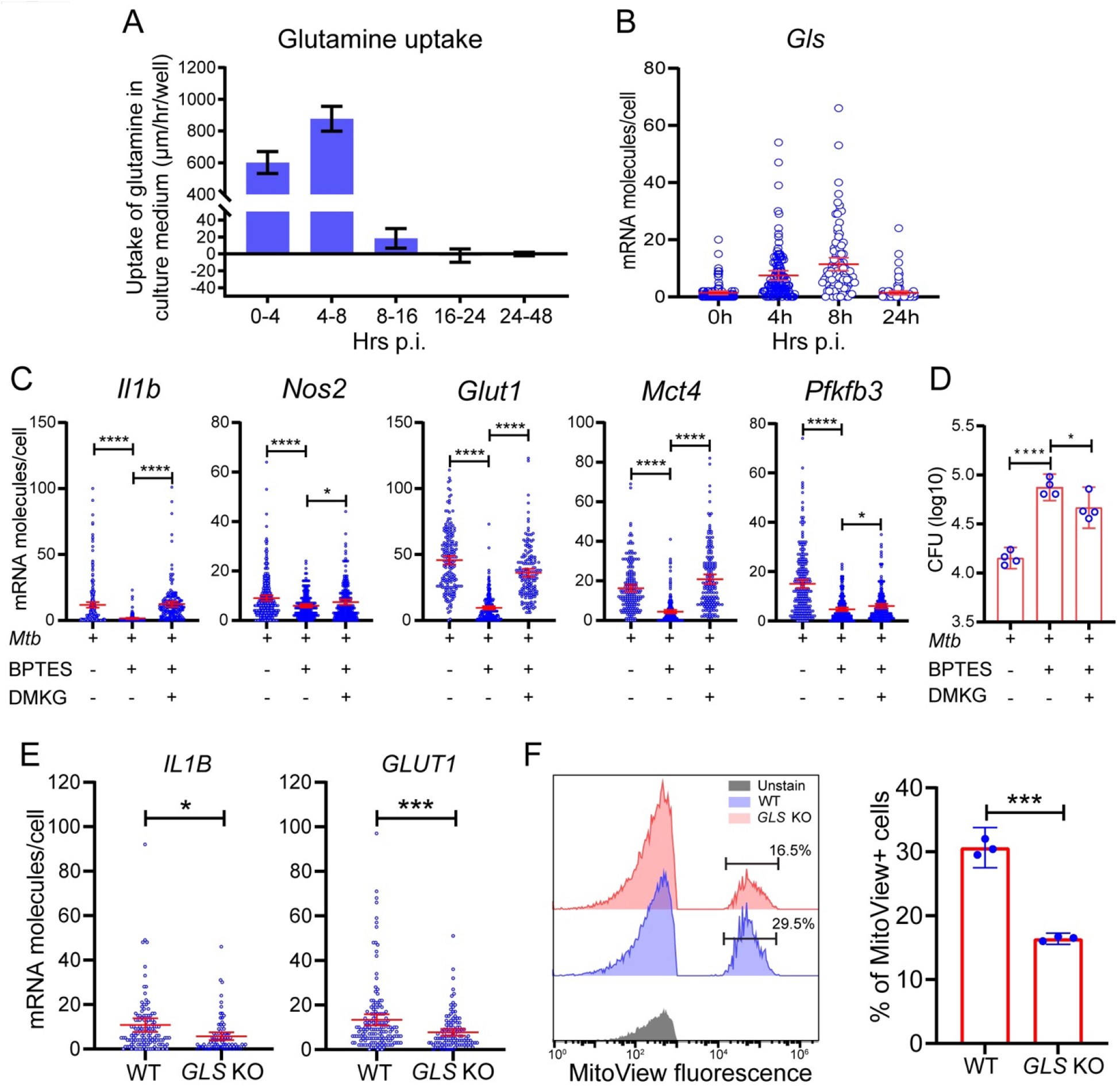
Requirement of glutamine for M1-like polarization. BMDMs were infected with *Mtb*, and cells and supernatants were collected at different times for single-cell mRNA analysis by sm-RNA-FISH and for measurement of glutamine uptake/utilization, respectively. Glutaminase (GLS) inhibitor BPTES was added to sets of cultures for GLS inhibition experiments at a final concentration of 10 μM. For rescue experiments, 1.5 mM dimethyl α-ketoglutarate (DMKG) was added to sets of cultures treated with the inhibitor. High rate of glutamine uptake/utilization by *Mtb*-infected macrophages corresponding to the M1-like polarization (**A**). Cell culture supernatants collected at the indicated times were subjected to glutamine determination using the Glutamine/Glutamate-Glo (TM) Assay kit (Promega, Madison, WI). Kinetics of glutamine uptake/utilization (μm per hour per well) were calculated based on its changes in the culture medium. Data are shown as means ± SDs from three independent experiments. Increased mRNA molecules for glutaminase gene *Gls* in M1-like macrophages (**B**). *Gls* mRNA molecules in infected BMDMs were detected and analyzed by sm-RNA-FISH as in Fig. 1. Diminished M1-like polarization by GLS inhibition with BPTES, and alleviation of the inhibition by treatment with DMKG (**C**). mRNA molecules in infected BMDMs, with or without 10 μM BPTES and/or 1.5 mM of DMKG at 8 hrs p.i were analyzed for *Il1b, Nos2, Glut1, Mct4*, and *Pfkfb3* by sm-RNA-FISH as in Fig. 1. Enhanced *Mtb* growth by GLS inhibition with BPTES (**D**). CFU of *Mtb* was determined by plating assay of cell lysates of infected BMDMs with indicated treatments at day 3 p.i. Dampened M1 polarization in THP-1 *GLS* KO cells (**E** - **F**). Wild-type THP1 cells, and *GLS* KO cells, generated by the Synthego Corporation (Menlo Park, CA), were subjected to differentiation and *Mtb* infection. mRNA expression of *IL1B* and *GLUT1* were analyzed by sm-RNA-FISH (**E**) as in Fig. 1. FISH data are shown as means ± 95% CI and represent three independent experiments. Mitochondrial mass was evaluated using MitoView Fix 640 (Biotium) by flow cytometry (left) and quantified (right) (**F**). Data are shown as means ± SDs from three independent experiments. Statistical significance at \**P* < 0.05, \*\**P* < 0.01, \*\*\**P* < 0.001 and \*\*\*\**P* < 0.0001 was based on two-tailed Student’s T-test.

We then used GLS inhibitors BPTES and CB-839 (51–53) to validate the role of glutamine catabolism in M1-like polarization. Treatment of infected BMDMs with BPTES or CB-839 dampened M1-like polarization, as measured by decreased mRNA levels of M1 immunometabolic markers that included *Il1b, Nos2, Glut1, Pfkfb3*, and *Mct4* (**Fig. 4C & S. Fig. 4**). The dampening consequently led to enhanced growth of *Mtb* in host cells (**Fig. 4D**). The inhibitory effects on macrophage polarization were alleviated by addition of dimethyl-α-ketoglutarate (DMKG), a membrane-permeable α-KG precursor (**Fig. 4C - D & S. Fig. 4**).

As a test for the generality of our observations with murine BMDMs, we examined the expression of *IL1B* and *GLUT1* in *Mtb*-infected THP1 cells in which *GLS* was knocked out (KO) using sm-RNA-FISH and mitochondrial mass staining with MitoView. At 8 hrs p.i., mRNA levels of *IL1B* and *GLUT1* and the mitochondrial mass were lower in KO cells that are deficient in *GLS* (**Fig. 4E - F**). Since an increase in mitochondrial mass is associated with macrophage proinflammatory differentiation and response (54), the decreased mitochondrial mass suggests a compromised proinflammatory differentiation in *GLS* KO cells. These data demonstrate the importance of glutaminolysis in the proinflammatory response of macrophages during *Mtb* infection.

## Conclusion

We conclude that glucose and glutamine play distinctive roles in the metabolic reprogramming during macrophage activation to the M1-like phenotype (**Fig. 5**). Besides the known role of increased glycolysis from glucose catabolism in promoting the M1-like polarization, our study identifies glutamine catabolism as an integral component of the metabolic reprogramming of M1-like macrophages by serving as an important source of carbon and nitrogen. A major role of glutamine catabolism during the M1-like polarization is to replenish the TCA cycle for the generation of TCA-cycle intermediates as signaling molecules, such as succinate which is responsible for HIF-1 stabilization and the switch of glucose metabolism to glycolysis (13, 55, 56), as well as biosynthetic precursors for the synthesis of aspartate and itaconate from OAA and aconitate, respectively. In addition, glutamine-derived glutamate and aspartate can potentially coordinate the metabolism of M1-like macrophages in different subcellular compartments. For example, glutamate participates in the redox homeostasis of M1 macrophages through the synthesis of GSH directly by serving as a substrate and indirectly by coupling with antiporter xCT for the uptake of cystine, another precursor for GSH synthesis (38, 57). Additionally, aspartate can partake in arginine regeneration in the cytosol in conjunction with NOS2-derived citrulline via the argininosuccinate synthase 1 (ASS1) and argininosuccinate lyase (ASL) pathways for sustained NO production by NOS2 of M1 macrophages (**Fig. 5**), especially when extracellular arginine levels are low (58). Furthermore, aspartate, together with glutamine, participates in the synthesis of nucleotides required for M1 polarization, probably through the purine nucleotide cycle (47). Finally, given that glutamate and aspartate are produced predominately in mitochondria by GLS and transamination reactions, respectively, a metabolic coordination between mitochondria and cytosol may constitute an important feature of the metabolic reprogramming of M1-like macrophages. Such coordination between the two compartments can be achieved by coupling two mitochondrial membrane transporters: the mitochondrial membrane glutamate carrier 1 (SLC25A22/GC1) and the aspartate-glutamate antiporter (AGC1/SLC25A12) (59) (**Fig. 5**). Gene expression for both transporters is increased in *Mtb*-infected M1-like macrophages (49). Indeed, the role of glutamine-derived aspartate in supporting cell growth and cellular redox homeostasis is supported by the observation that insufficient cytosolic aspartate delivery leads to cell death when TCA-cycle carbon is reduced following glutamine withdrawal and/or glutaminase inhibition by the small-molecule inhibitor CB-839 (60). Collectively, the pleiotropic routes of glutamine catabolism linking metabolic pathways between cytosol and mitochondria during the M1-like polarization indicate an intimate metabolic coordination of intracellular processes to meet the requirement of M1-like macrophages for bioenergetics and biosynthetic precursors (**Fig. 5**), as is seen in proliferating cells (61, 62).

**Fig. 5.**
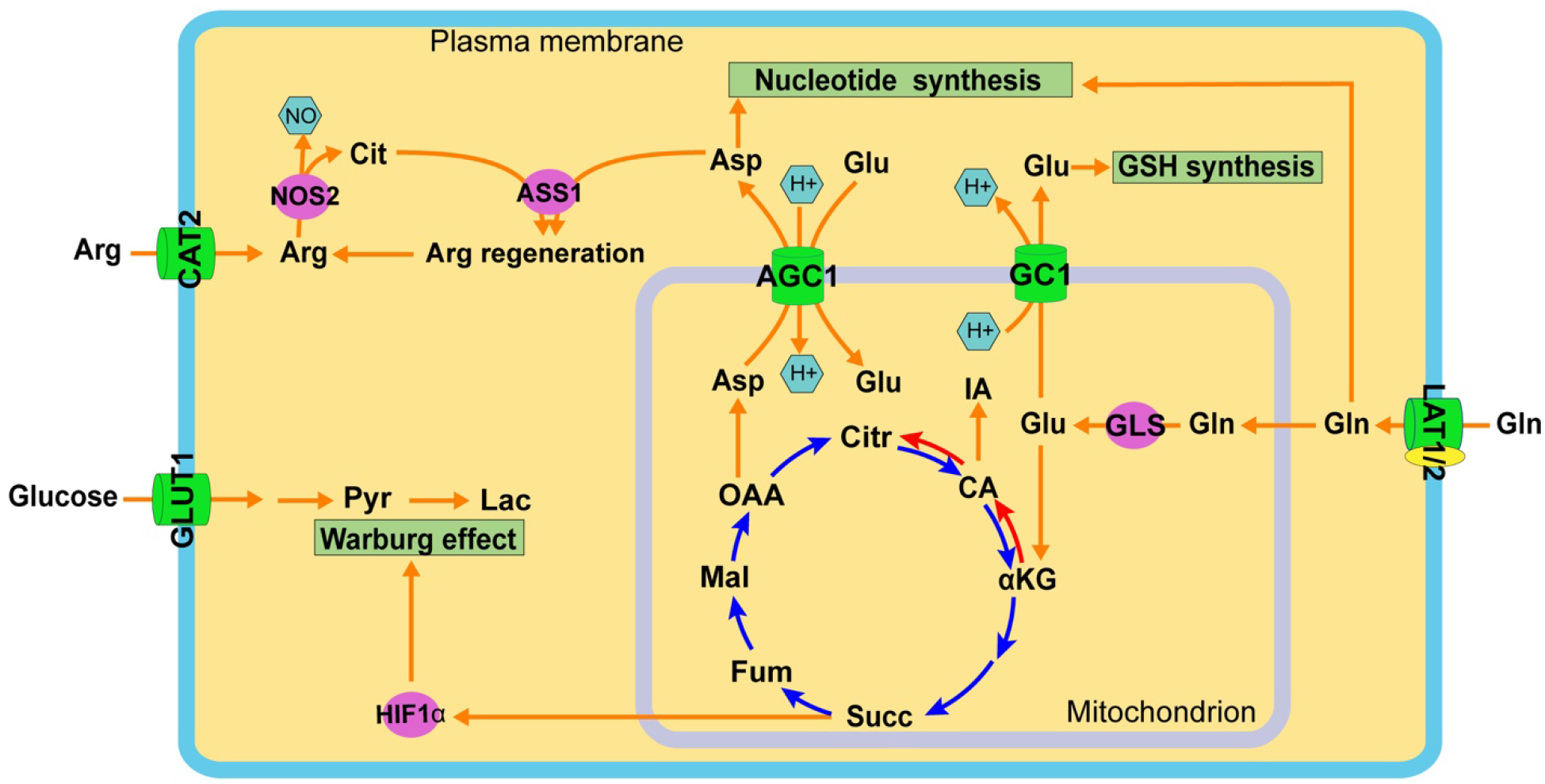
Coordination of glucose and glutamine catabolism during M1-like polarization. M1-like polarization is marked with increased catabolism of glucose and glutamine (Gln) in distinctive subcellular compartments, but with coordinated functions. Increased glucose uptake, mediated by GLUT1, and its catabolism in the glycolysis pathway in cytosol result in the production of lactate (Lac). Increased Gln uptake, probably mediated by neutral amino acid antiporters LAT1 and/or LAT2, and its direct metabolite glutamate (Glu) participate in multiple pathways of the metabolic remodeling program of M1-like macrophages. Glu, the product of the glutaminolysis pathway by glutaminase (GLS) in mitochondria, enters the TCA cycle for anaplerosis reactions. Glu replenishes the TCA cycle leading to the generation of the antimicrobial metabolite itaconate (IA) and the signaling molecule succinate, which promotes stabilization of HIF1α and the subsequent Warburg Effect. Gln also contributes both carbon and nitrogen to the formation of the nonessential amino acid aspartate (Asp), probably as a result of transamino reactions in mitochondria, which is then involved in biosynthetic pathways in the cytosol, including nucleotide synthesis and intracellular arginine regeneration by coupling with nitric oxide synthase 2 (NOS2)-derived citrulline (Cit) via argininosuccinate synthase 1 (ASS1) to sustain NO generation by NOS2. The export of mitochondrial Asp to the cytosol is mediated by the functional coupling between two mitochondrial carriers, wherein the glutamate carrier (GC1) exports the mitochondrial Glu deriving from GLS activity to cytosol. Cytosolic Glu then serves as a substrate of aspartate/glutamate carrier 1 (AGC1) for the net export of mitochondrial aspartate to the cytosol. Cytosolic Glu also participates in intracellular redox homeostasis involving the synthesis of glutathione (GSH). Abbreviations: CAT2: cationic amino acid transporter 2; αKG: α-ketoglutarate; Arg: Arginine; CA: cis-aconitate; Citr: citrate; Fum: fumarate; Mal: malate; OAA: oxaloacetate; Pyr: pyruvate.

Our study has potential limitations. For example, the data were derived from a snapshot of the metabolic state of M-1 like polarization during *Mtb* infection, and they do not necessarily represent the metabolic steady-state of the M1-like macrophages. In addition, the detection limitation and sensitivity of the isotope analytical systems, together with possible low concentrations of some metabolites, could be responsible for the absence of ^13^C enrichment for key metabolites such as α-KG. Nevertheless, the overall metabolomics data clearly demonstrate that glutamine contributes carbon and nitrogen to multiple cellular processes of the metabolic reprogramming that occurs during *Mtb* infection-induced M1-like polarization (**Fig. 5**). Since findings from cytokine quantitative trait loci analysis show that glutamine metabolism-related genes are associated with the cytokine response of peripheral blood cells to *Mtb* infection (63), glutamine likely plays critical roles in host immunity against *Mtb* infection *in vivo*. The present study is a first step toward understanding the role of glutamine metabolism in different pathophysiological states of TB disease and will guide the development of host-directed therapies to improve bacterial clearance and/or prevent the induction of immunopathology.

## Material and Methods

### *Mtb* culture

Cultures of GFP-*Mtb* H37Rv were grown under aerobic conditions at 37°C in Dubos Tween Albumin (DTA) medium (Becton Dickinson, Franklin Lakes, NJ) (64). Mid-log phase culture at an OD of 0.3 - 0.5 at 580 nm was used for infection.

### Culture, differentiation, and infection of mouse BMDMs and THP1 cells

Bone marrow cells isolated from femur and tibia of C57BL/6J mice were differentiated into BMDMs for 7 – 8 days by a standard procedure (65) in DMEM (Sigma, St. Louis, MO) supplemented with 10% fetal bovine serum (FBS) (Thermo Fisher, Waltham, MA), 10% L929-cell-conditioning media, and 1% penicillin/streptomycin (Corning, Glendale, AZ) (49). BMDMs were seeded into 6- and 24-well plates at a density of 1X10^6^ and 3X10^5^ cells per well, respectively. For sm-RNA-FISH, BMDMs were seeded onto gelatin-coated coverslips located at the bottom of 24-well plates. BMDMs were infected with *Mtb* at a multiplicity of infection (MOI) of 4 in culture medium for 4 hrs, and they were further cultured for up to 24 hrs after the removal of extracellular bacteria. For GLS inhibition experiments, inhibitors CB-839 or BPTES (MedChemExpress LLC, Monmouth Junction, NJ) were added to sets of cultures 12 hrs prior to infection at a final concentration of 10 μM, according to literature (66, 67). For rescue experiments, dimethyl α-ketoglutarate (DMKG, Sigma, St. Louis, MO) was added to sets of cultures with either inhibitor at a final concentration 1.5 mM (68). After removal of extracellular bacteria at 4 hrs p.i, fresh culture media containing same concentration of either inhibitor or DMKG were added back to the corresponding wells for the duration of the experiments. BMDMs and supernatants were harvested at different times p.i. together with those from uninfected controls for various analyses. For *Mtb* CFU determination in macrophages, an MOI of 1 was used for infection, cells were lysed in 0.6% SDS solution at the indicated times, and CFU was determined by plating cell lysates on 7H11 agar plates following standard procedures (64).

THP1 cells (ATCC: TIB-202, Manassas, VA) were used to generate a pool of *GLS*-null cells. CRISPR/Cas9-mediated knockout (KO) cells were generated by the Synthego Corporation (Menlo Park, CA). *GLS* KO cells and their parental wild-type cells were cultured in RPMI cell culture medium (Sigma, St. Louis, MO) supplemented with 10% FBS (Thermo Fisher, Waltham, MA), differentiated by treatment with 100 μM of phorbol 12-myristate 13-acetate (PMA, Sigma, St. Louis, MO) for 24 hrs, recovered for 24 hrs in fresh RPMI medium, and then infected as described for mouse BMDMs.

### sm-RNA-FISH

sm-RNA-FISH probes were designed using Stellaris™ probe designer, an online tool (https://www.biosearchtech.com/products/rna-fish). About 50 3’-amino modified oligonucleotides for each mRNA transcript were obtained from LGC Biosearch (Petaluma, CA) (the probe sequences will be provided upon request). Pooled oligonucleotides were coupled with tetramethylrhodamine (TMR), Texas Red, or Cy5 fluorophores and purified by HPLC, as detailed in (69). BMDMs at different times of infection were fixed by 10% formaldehyde and permeabilized in 70% ethanol. After equilibration in hybridization wash buffer (10% formamide (Thermo Fisher, Waltham, MA) in 2X SSC), cells were incubated overnight in hybridization buffer (10% formamide (Thermo Fisher, Waltham, MA), 10% dextran sulfate (Sigma, St. Louis, MO), 2 mM vanadyl-ribonucleoside complex (Sigma, St. Louis, MO), 0.02% RNAse-free BSA (Thermo Fisher, Waltham, MA), 0.001% *E. coli* tRNA (Sigma, St. Louis, MO) containing labeled mRNA probes at 37°C. The pooled probe set for each target mRNA was used at 25 ng per hybridization reaction (50 μl). Following incubation, cells were washed in hybridization wash buffer followed by mounting with antifade buffer (Abcam, Cambridge, UK) before proceeding to imaging.

### Image acquisition and analysis

Analysis of mRNA molecules was performed using a modified system developed by Raj, et al (69). Briefly, coverslips with stained cells were placed on an Axiovert 200M inverted fluorescence microscope (Zeiss, Oberkochen, Germany) with a 63x oil-immersion objective (numerical aperture 1.4), a 14 Prime sCMOS camera (both from Photometrics, Tucson, AZ), and the system was controlled by Metamorph image acquisition software (Molecular Devices, San Jose, CA). Images in DIC, GFP, and each of the other fluorescence channels corresponding to the fluorophores used in each probe set. Images from fluorescence channels consisted of 16 optical sections separated by 0.2 μm with an exposure time ranging from 1500-2,000-milliseconds. Image analyses were performed using modified image processing programs in MATLAB R2021a (Natick, MA). Briefly, the boundaries of all cells in each field were first charted by manually tracing over their DIC image to avoid biases. Z-stacks of RNA images were then analyzed, and the number of spots corresponding to RNA molecules in each fluorescence channel was counted. The algorithm processes the images with a Laplacian filter and provides the user with a 3D plot for a region of the cells with spots, allowing users to provide a threshold to separate the noise from the signal. This threshold was then employed to calculate the number of spots within the boundaries of cells. Evidence for the accuracy of this algorithm in counting cellular RNA molecules has been published (27, 69).

### Profiling of widely targeted small metabolites by QTRAP 6500+ LC-MS/MS Systems (ABSciex)

BMDMs obtained by standard procedure were seeded into 6-well plates (1X10^6^ cells per well) in DMEM with 10% dialyzed FBS (Thermo Scientific, Waltham, MA), 4 mM glutamine, and 25 mM glucose but without pyruvate one day prior to infection. BMDMs were infected by *Mtb* in the above-described DMEM medium for 4 hrs and cultured for another 4 hrs after the removal of extracellular bacteria. Cells at 8 hrs p.i and the corresponding uninfected control cells were harvested by centrifugation, and metabolites were extracted with 80% methanol with internal standards of small metabolites by multiple rounds of freeze-and-thaw cycle in liquid nitrogen and ultrasonic ice water bath, respectively. Metabolites in extraction solvent was collected by centrifugation and dried under gentle nitrogen flow. Metabolites were reconstitute in 80% methanol and analyzed with UPLC coupled with ABSciex 6500+ QTrap mass spectrometer (ABSciex, Framingham, MA). Briefly, metabolite separation was performed on a reverse phase ACE PFP-C18 column, and data were collected with a Multiple Reaction Monitoring (MRM) mode, using MultiQuant™ software (Sciex, Framingham, MA) to enable the identification and quantification of metabolites of interest. A pooled quality-control sample was injected six times and used to calculate the coefficients of variation (CV). Metabolites with CV higher than 30% were excluded from the data analysis. Multivariate analysis of the data set was performed by SIMCA-p software (Sartorius, Goettingen Germany), and the partial least squares-discriminant analysis (PLS-DA) model was used to demonstrate differences in the metabolic profiles between infected macrophages and uninfected controls. Analysis of pathway enrichment was carried out by Metaboanalyst (V5.0) (https://www.metaboanalyst.ca).

### Tracing metabolomics by U^13^C glutamine, U^15^N glutamine, and U^13^C glucose

BMDMs were seeded into 6-well plates (one million cells per well) in DMEM containing 10% dialyzed FBS (Thermo Scientific, Waltham, MA), 4 mM unlabeled glutamine, and 25 mM glucose one day before infection. One hour before the infection, the culture medium was replaced by DMEM containing 10% dialyzed FBS, but without glutamine. BMDMs were infected by *Mtb* in DMEM medium supplemented with 10% dialyzed FBS and 4 mM of 50% U^13^C glutamine (Cambridge Isotope Laboratories, Inc, Tewksbury, MA) (2 mM labeled and 2 mM unlabeled glutamine) for 4 hrs. After removing extracellular bacteria cells were cultured for another 4 hrs in DMEM medium supplemented with 10% dialyzed FBS and 4 mM of 50% U^13^C glutamine. For tracing experiments with U^15^N glutamine, 4 mM of 50% of U^15^N glutamine (2 mM labeled and 2mM unlabeled glutamine in replacement of U^13^C glutamine) was applied to the cultures.

For tracing experiments with U^13^C glucose, BMDMs were seeded into 6-well plates (one million per well) in DMEM with 10% dialyzed FBS (Thermo Scientific, Waltham, MA), 25 mM unlabeled glucose, and 4 mM glutamine one day before the infection. One hr before infection, the culture medium was replaced by DMEM with 10% dialyzed FBS (Thermo Scientific, Waltham, MA) but lacking glucose. BMDMs were infected by *Mtb* in DMEM medium supplemented with 10% dialyzed FBS and 25 mM of 50% U^13^C glucose (Cambridge Isotope Laboratories, Inc, Tewksbury, MA) (12.5 mM labeled and 12.5 mM unlabeled glucose) for 4 hrs. After extracellular *Mtb* removal, cells were cultured for another 4 hrs in DMEM medium supplemented with 10% dialyzed FBS and 25 mM of 50% U^13^C glucose.

Uninfected control sets under the respective culture conditions with 4 mM 50% of U^13^C glutamine, U^15^N glutamine, or 25 mM 50% of U^13^C glucose were included for comparative analyses. Natural abundance subtraction samples were also prepared with unlabeled 4 mM glutamine or 25 mM glucose. Cells at 8 hrs p.i. and corresponding uninfected controls were detached and harvested by centrifugation. Metabolite extraction from cell pellets was performed with 80% methanol as described above. The extract was dried under gentle nitrogen flow, and derivatized with a methyl-moximation (with 15 mg/ml methoxyamine in pyridine, 30 ^0^C for 90 min) and silylation (BSTFA or MTBSTFA, at 70 ^0^C for 60 min). The samples were then analyzed with GC-TOF/MS (Waters, Milford, MA) with an electron impact mode and a DB-5MS column (Agilent, Santa Clara, CA) following our published protocol (39). The data were analyzed with Masslynx software (Waters, Milford, MA), and the enrichment calculation followed Jennings and Matthews (70).

### Glutamine Measurement

Culture supernatants from *Mtb*-infected BMDMs at 0, 4, 8, 16, 24, and 48 hrs p.i were collected by filtering with 0.2 μM microcolumn by centrifugation. Glutamine in culture supernatant was determined using the Glutamine/Glutamate-Glo (TM) Assay kit (Promega, Madison, WI) and used for calculating the kinetics of glutamine uptake/utilization, following manufacturer’s instructions.

### ELISA

Culture supernatants from *Mtb*-infected BMDMs at 0, 4, 8, and 24 hrs p.i were collected by filtering with 0.2 μM microcolumn by centrifugation, and then subjected to measurement of IL-1β protein using the Mouse IL-1 beta-uncoated ELISA kit (Invitrogen, Waltham, MA), following manufacturer’s instructions.

### Flow cytometry

Infected THP1 cells were stained with 100 nM MitoView FIX 640 (Biotium, Fremont, CA) at the indicated time points. Cells were then fixed in 4% formaldehyde, collected, and analyzed on a BD FACS Celesta (Becton Dickinson, Franklin Lakes, NJ) to evaluate mitochondrial mass. Data were analyzed using FlowJo version 7.5.5 software (Tree Star Inc., Ashland, OR).

### Statistics

A 95% confidence interval (CI) and/or two-tailed Student’s T-test among groups were carried out for statistical significance analyses. Multivariant statistical analysis for the widely targeted metabolite data was performed with SIMCA-p software (Sartorius, Goettingen Germany). PLS-DA model was performed with unit variant scaled data. The cut-off for the variable importance in the projection (VIP) value was set to 1. Pathway analysis was performed in Metaboanalyst (V5.0) (https://www.metaboanalyst.ca). Comparison of isotope distribution in metabolites between infected cells and uninfected controls was analyzed by two-tailed Student’s T-test using GraphPad Prism 8.0 (San Diego, CA).

## Declarations

### Consent for publication

All authors agree with the publication.

### Availability of data and materials

yes.

## Acknowledgment

We thank Jason Yang and Ryan Dikdan for critical reading of the manuscript.

## Authors’ contributions

LS and QJ conceived the concept, and LS, QJ, YQ, IJK and ST designed the experiments. QJ, YQ and LS performed experiments, and QJ, LS, YQ, IJK and ST analyzed data. LS, QJ, YQ, IJK, SS, and KD drafted the manuscript and QJ and YQ prepared the figures. All authors contributed to the article and approved the submission.

